# Spatiotemporal Decoding of Explore-Exploit Decisions in the Human Brain

**DOI:** 10.64898/2026.06.01.729427

**Authors:** Rohit Yadav, John D. Romero, Julia M. Stephen, Jon M. Houck, James F. Cavanagh, Vincent D. Costa, Jeremy Hogeveen

**Affiliations:** Psychology Clinical Neuroscience Center; Department of Psychology, University of New Mexico; University of Washington, Seattle, Washington; The Mind Research Network, a Division of Lovelace Biomedical Research Institute; Department of Psychiatry and Behavioral Sciences, Emory University School of Medicine, Atlanta, GA 30329

**Author notes:** Manuscript correspondence: Dr. Rohit Yadav or Dr. Jeremy Hogeveen.

**Keywords:** Explore-Exploit, Value-Based Decision-Making, Prefrontal Cortex, Magnetoencephalography (MEG), Reinforcement Learning

## Abstract

Adaptive behavior requires flexibly shifting between exploiting familiar rewards and exploring novel opportunities. These explore-exploit decisions are implemented via a distributed brain network—anchored in frontopolar cortex (FPC) and ventromedial prefrontal cortex (vmPFC)—that computes the total value of a given choice by weighting the immediate value of familiar options against the latent future value of exploration. Capturing the precise temporal dynamics of these neural computations across spatially distributed cortical networks is critical to resolving how the brain manages this tradeoff. Here, we combined magnetoencephalography (MEG) with partially observable Markov decision process (POMDP) modeling during a reinforcement learning task. By decoding POMDP-derived choice policies in MEG source space, we mapped the precise spatiotemporal emergence of explore-exploit decisions across the cortex. We demonstrate that explore-exploit policy implementation unfolds via a hierarchical functional dissociation across the rostral prefrontal cortex. During decision formation, the lateral FPC initiates the strategic shift toward exploration hundreds of milliseconds before choice execution. Conversely, vmPFC and orbitofrontal cortex (OFC) do not exhibit early significant divergence from the baseline exploitative trace on exploration trials, displaying only a delayed, transient peak prior to choice execution. Following feedback, the vmPFC and OFC transition to a sustained representation of the choice’s empirical value to update reward expectancies and guide future actions. Mapping reinforcement learning variables onto millisecond-resolved neural data reveals how the human brain resolves the explore-exploit dilemma: through early strategic initiation of exploration in the FPC, a delayed exploratory shift in vmPFC and OFC, and sustained outcome evaluation to optimize future actions.

## INTRODUCTION

Adaptive decision-making requires an optimal balance between exploiting familiar rewards and exploring novel opportunities. Managing these explore-exploit tradeoffs requires the brain to compute and weigh two distinct types of information: the learned value of familiar choice options against the motivational salience of trying something new (Averbeck, 2015; Frank et al., 2009; Wilson et al., 2014, 2021). While frameworks for studying this tradeoff vary across the literature, a consensus across human and nonhuman primate computational neuroimaging is that these choice computations are reliably encoded within a set of prefrontal decision-making hubs. Specifically, ventromedial prefrontal and orbitofrontal cortices (vmPFC and OFC, respectively) are involved in learning to exploit familiar options and tracking expected value estimates (Blanchard & Gershman, 2018; Cockburn et al., 2022; Costa & Averbeck, 2020; Daw et al., 2006; Hogeveen et al., 2022; Wyatt et al., 2024). In contrast, the lateral frontopolar cortex (FPC) is thought to compute the informational value of state uncertainty to drive directed exploration (Badre et al., 2012; Cockburn et al., 2022; Daw et al., 2006; Hogeveen et al., 2022; Zajkowski et al., 2017). Additionally, behavioral shifts between exploration and exploitation appear to rely on the cingulo-opercular network (Blanchard & Gershman, 2018; Sazhin et al., 2025), as well as subcortical structures like the amygdala and striatum (Costa et al., 2019). However, focusing primarily on *where* these decision policies are encoded has limited our understanding of *how* these computations emerge in real-time across distributed brain networks.

Emerging circuit-level accounts propose that explore-exploit decisions may actually emerge from rapid, brain-wide cascades of motivationally relevant information. Previously, we used model-based fMRI with a whole-brain focus to demonstrate that the latent future value of exploration is not just encoded in frontopolar and ventromedial prefrontal regions, but recruits a brain-wide network including several previously underappreciated parietal, temporal, and occipital regions (Hogeveen et al., 2022). Studies using magnetoencephalography (MEG), which blends the temporal resolution of EEG with spatial resolution comparable to fMRI, suggest that rather than a monolithic prefrontal computation, value-based and explore-exploit decisions rely on highly coordinated, time-varying cortical hierarchies to initiate, execute, and monitor ongoing choice policies (Hallquist et al., 2024; Hunt et al., 2012, 2015). Consequently, a fundamental tension has emerged in decision neuroscience: Are the empirical and latent value computations underlying explore-exploit decision-making computed within isolated prefrontal decision-making modules, or are they an emergent property of a dynamic, cortex-wide hierarchical cascade of neural computations? Resolving these competing frameworks requires mapping the precise spatiotemporal architecture of the computational signals involved in managing explore-exploit tradeoffs.

Crucially, these two conceptual accounts are not mutually exclusive; decision-making may emerge from a dynamic cortical flow that ultimately converges to resolve a goal-directed choice policy within the prefrontal cortex. To test this unified spatiotemporal account, we recorded MEG during a novelty-bandit reinforcement learning task, operationalizing choice computations via a Partially Observable Markov Decision Process (POMDP) framework that formally quantifies both the immediate expected value (IEV) of exploitation and the latent future value of exploration (BONUS). By decoding choice policies in source space, we demonstrate that explore-exploit decisions do not emerge instantaneously via a monolithic prefrontal computation. Instead, the lateral frontopolar cortex (FPC) initiates the strategic shift toward exploration early in the decision window, whereas the ventromedial prefrontal (vmPFC) and orbitofrontal cortex (OFC) exhibit only delayed, transient exploratory modulations prior to choice execution. Upon receiving feedback, this network functionally reconfigures, with the vmPFC and OFC transitioning to a sustained encoding of empirical choice value. Ultimately, by mapping computational variables onto millisecond-resolved neural data, we provide direct evidence that navigating the explore-exploit dilemma relies on a dynamic sequence of distinct prefrontal operations.

## METHODS AND MATERIALS

### Human Participants

28 healthy adult participants (15 females, 13 males; M = 26.6 years, SD = 7.24 years) recruited from the Albuquerque, New Mexico community were included in the final analyses. All experimental procedures were approved by the University of New Mexico Office of the Institutional Review Board, and all participants provided written informed consent before participating. Data from an additional 2 participants were excluded prior to analysis due to excessive neuroimaging artifacts, insufficient task response rates, or extreme behavioral anomalies relative to the computational model.

### Novelty-Bandit Task

A schematic of the task design is included in **Figure 1.A**. Participants completed a speeded choice task in which they had up to 2 seconds to make manual responses between three neutral images drawn from the International Affective Picture System (IAPS) (Bradley & Lang, 2020). Stimuli were presented using E-Prime Version 3 (Psychology Software Tools, Sharpsburg, PA), and responses were recorded with the MIND Input Device^1^. At the start of each set, the three images were randomly assigned reward probabilities of low (p = 0.2), medium (p = 0.5), or high (p = 0.8). Sets remained stable for 5–12 trials (M = 7 ± 2). After this interval, insertion of a novel choice option occurred. One of the familiar images from the prior set was replaced with a novel image. This manipulation introduced uncertainty about the assigned reward probability of the novel option while making the explore-exploit tradeoff explicit. Each novel image was randomly assigned a reward probability from the set of three possible reward probabilities, with the constraint that no set containing three images was assigned the same reward probability (i.e. a preferred option could always be identified). On each trial, we randomized the spatial position of the images, and re-mapped the response buttons accordingly (i.e., index, middle, and ring finger presses corresponded to the left, middle, and right spatial positions, respectively). Following each choice, participants received immediate feedback indicating either reward (a green “+1”) or no reward (a red “0”). Participants also reported their confidence (low, medium, or high) after each choice. These confidence ratings are not analyzed in the present manuscript.

**Figure 1.**
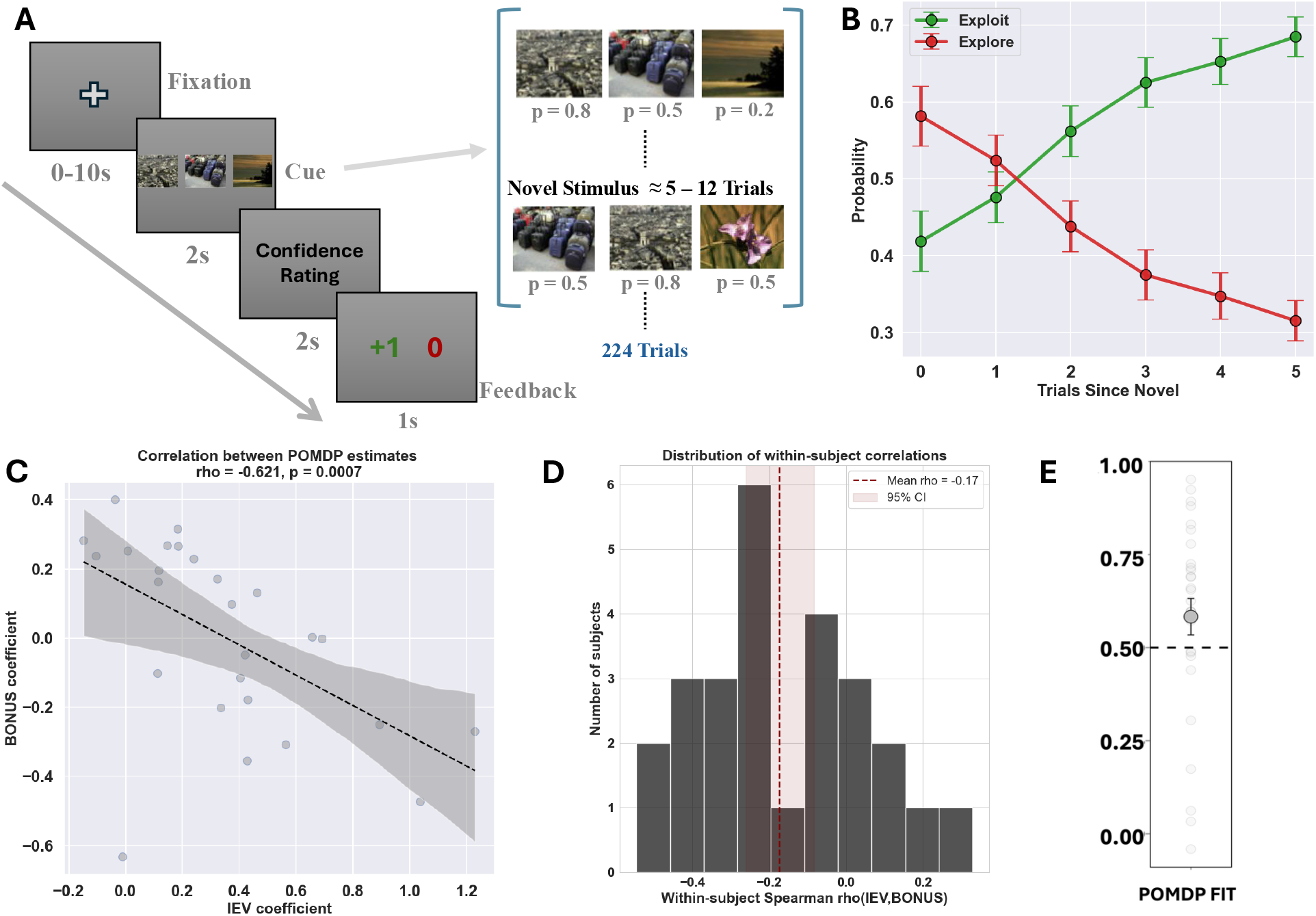
Task design, behavioral performance and computational model. (A) Trial events from the three-arm bandit tasks and schematics (cf., Hogeveen et al., 2022). (B) Participants initially preferred exploratory over exploitative choices following the introduction of novel stimuli, consistent with adaptive responses to uncertainty. (C) The parameter estimates used to weight IEV, and exploration BONUS were negatively associated. (D) Histogram of the correlation between BONUS and IEV parameters within subjects (i.e., trial-level estimates). Overall, there was a small negative effect size for this relationship across the sample. (E) Choice performance was strongly correlated with the POMDP model (> 0.5), indicating that participants explore–exploit behavior aligns with the computations captured by the model.

### Computational Modeling

To characterize the computational processes underlying behavior, choices were modeled using a partially observable Markov decision process (POMDP), that estimates the utility of selecting each option while accounting for uncertainty about future outcomes (Averbeck, 2015). In this framework, the utility of an option is defined as the sum of two components: the immediate expected value (IEV) and the future expected value (FEV). IEV reflects the probability that selecting an option will result in an immediate reward and can be estimated from the history of observed outcomes. FEV reflects the discounted value of future rewards expected following a given choice and captures the potential long-term benefit of sampling an option. To quantify the value of exploration, we computed an exploration bonus (BONUS), defined as the difference between the FEV of a given option and the average FEV across all available options. Positive BONUS values indicate that sampling an option may provide greater future information or reward potential, whereas negative BONUS values indicate that the option is relatively well characterized. Exploration BONUS and IEV estimates were computed for all three options on a trial-by-trial basis. In the present task, novel options were introduced periodically. Because their reward probability was initially uncertain, newly introduced options were associated with high BONUS values that decreased as the option was sampled over successive trials. Under this framework, choices associated with positive BONUS values were considered exploratory, whereas choices with negative BONUS values were considered exploitative.

### MEG Sensor-level processing

MEG data were acquired using a 306-channel MEGIN Neuromag system (Elekta AB, Stockholm, Sweden), comprising 102 magnetometers and 204 planar gradiometers. Data were recorded continuously at a sampling rate of 1000 Hz. All preprocessing was carried out using MNE-Python (Gramfort et al., 2013). Muscle artifacts were detected using a z-score–based method from MNE applied to high-frequency activity (110–140 Hz), and contaminated segments were annotated. Head motion was estimated from continuous head position indicator (cHPI) signals, and time periods exceeding a displacement threshold of 2 cm were annotated; all annotated segments were subsequently dropped. Noisy and flat channels were automatically identified using the Maxwell filtering-based algorithm (Taulu & Hari, 2009) and removed. Signal Space Separation (SSS) with temporal extension (tSSS) was then applied using a maxwell filter to suppress external noise and compensate for head movement. Data were transformed to a common head position and corrected using calibration and crosstalk files. For epoching, signal-space projection (SSP) vectors derived from electrooculogram (EOG), and electrocardiogram (ECG) recordings were applied to remove ocular and cardiac artifacts. The data were subsequently low pass filtered at 40 Hz and downsampled to 250 Hz. Epochs were extracted at cue onset events (−200 to 1200ms) and at feedback onset (−200 to 1000ms). A pre-stimulus baseline correction was applied to each epoch using the 200ms interval preceding event onset, whereby the mean signal during this window was subtracted from the entire epoch to remove slow drifts and offset differences. Trials without a valid behavioral response were excluded, and the resulting cleaned epochs were retained for subsequent sensor and source-level analyses.

### MEG Source-level processing

Source reconstruction was performed using MNE-Python (Gramfort et al., 2013) in conjunction with anatomical MRI data processed with FreeSurfer (Fischl, 2012). For each participant, a three-layer Boundary Element Model (BEM) was constructed from T1-weighted images using standard watershed segmentation, comprising inner skull, outer skull, and scalp compartments. MEG and MRI coordinate systems were aligned via rigid-body co-registration using fiducial landmarks and digitized head shape data with manual alignment. Source estimates were computed using a minimum-norm inverse solution (MNE) based on the BEM forward model. Noise covariance matrices were estimated from the pre-stimulus baseline (*t* = -200ms to 0ms) and incorporated into the inverse operator. The resulting estimates were noise-normalized using dynamic statistical parametric mapping (dSPM), with regularization determined by an assumed signal-to-noise ratio of 3 (λ^2^ = 1/SNR^2^). We note that activity estimated in subcallosal regions can be less reliable due to reduced sensitivity and spatial leakage inherent to MEG, and exclusion of these regions in visualization was performed to avoid overinterpretation of such effects. This masking was applied post hoc and did not affect the inverse solution, all cortical vertices were included during source estimation.

### Decoding and Encoding

#### Temporal and Spatiotemporal Decoders

To obtain a time-resolved measure of task-related information, temporal decoders were first used on sensor-level MEG activity. Multivariate pattern classification was implemented using the MNE-Python and scikit-learn framework (Gramfort et al., 2013; Pedregosa et al., 2012). At each time point, the pattern of activity across sensors was treated as a feature vector, and a logistic regression classifier with L2 regularization was trained to discriminate task conditions independently at each time point. Classifier performance was evaluated using 5-fold stratified cross-validation across trials. Decoding performance was quantified using the area under the receiver operating characteristic curve (AUC;(Fawcett, 2006)), providing a bias-free measure of classification accuracy. This approach provides a time-resolved estimate of task-relevant information, identifying the precise moments when decision strategies (at cue) and outcomes (at feedback) are reliably represented in sensor-level activity.

In contrast to sensor-level temporal decoding, source-space decoder was trained on patterns of activity across the entire set of cortical sources spanning the whole brain at each time point. The decoder was trained and evaluated independently at each time point, preserving the temporal resolution of the analysis. At the same time, it leverages the full spatial structure of source-space activity across the cortex, allowing the identification of informative regions and their contributions over time. To avoid cross-subject alignment bias and preserve individual structural variations, all localized features/ROIs were extracted within each participant’s native space.

#### Haufe Coefficients

To further enable neurophysiological interpretation of the spatial decoding models, linear decoder weights **W** were transformed into activation patterns **A**, using Haufe transformation (Haufe et al., 2014), with implementation provided in the MNE-Python toolbox (Gramfort et al., 2013). While raw decoder weights are optimized to suppress noise and maximize classification accuracy, they often exhibit spatial leakage, where weight is assigned to sources with no relevant neural signal but simply to cancel out correlated noise. Therefore, raw linear weights from the decoder reflect a combination of task-relevant signals and the covariance structure of the data that are not directly interpretable. By applying the Haufe transformation, these weights are re-scaled by the covariance of the MEG data **Σ**_**x**_, effectively re-projecting the information back into the measurement space using the transformation **A** = **Σ**_**x**_ **WΣ**_**s**_^**−1**^, where **Σ**_**s**_ the covariance of the target variable. This creates activation patterns that more directly reflect the contribution of neural sources to the decoded signal. As the sign of these patterns can still be influenced by arbitrary factors such as class labels, model regularization, and correlations between sources, we focus on their magnitude (absolute value) to quantify the strength of neural engagement. These transformed coefficients therefore allow for a robust comparison of neural engagement across our defined ROIs, ensuring that the spatiotemporal cascades identified in our results (**Figure 3**) reflect genuine physiological engagement obtained from decoding.

#### Univariate Encoders

To characterize the time-resolved encoding of decision strategies, a mass univariate regression framework was applied to the source reconstructed neural activity within regions of interest. At each vertex and time point (**t**), a general linear model was fitted to the single trial source data ***Y***_***i***_(***t***) = ***β***_**0**_(***t***) + ***β***_***cond***_(***t***) . ***X***_***i,cond***_ + ***ϵ*** , where ***Y***_***i***_(***t***) is the neural activity of a given vertex at time point (**t**) for trial **i, X**_**i,cond**_ represents the binary condition vector indicator (*Exploit = 0, Explore = 1*) for trial **i**, therefore ***β***_**0**_ represents the *exploit* baseline activity and ***β***_***cond***_ isolates the specific differential effect of *exploration*. Trial counts for the two conditions were balanced within each participant to avoid unbiased estimation by pooling a random subset of trials from the more frequent condition. To prevent the mathematical cancellation of opposite-signed regression coefficients due to individual variations in source geometry, we focused on effect magnitude; beta weights specific to each vertex were converted to absolute values and subsequently averaged across all vertices within each respective ROIs. This resulted in two simplified time courses per subject, capturing the overall magnitude of the exploit baseline and the explore deviation.

#### Decoder decision values

To track the time-resolved confidence and certainty of the neural representations from decoders, time-resolved trial-wise decision values (***z***(***t***)) were extracted from the multivariate decoding models. For each individual trial, the decision value was operationalized as the signed distance of the neural feature vector from the decoder’s optimal decision boundary at each timepoint. This relationship is reflected in the classification performance, where the rise in AUC corresponds to increasing distances from the decision boundary, indicating certainty level in the neural separability, providing a time-resolved index of how strongly neural activity patterns encode categorical information, where the logistic regression decision threshold is defined at 0, ***z***(***t***) = **w**(**t**)^**T**^**x**(**t**) + **b**(**t**), here **w** represents the vector of trained decoder weights, **x** is the neural activity pattern from sensor/source units at a given time point, and **b** is the intercept.

#### Statistical significance tests

A permutation-based cluster-correction was used to identify significant temporal clusters for both decoding (AUC) and encoding **(*β***) results using the implementation from MNE-Python (Gramfort et al., 2013). This test provides a principled way to control for multiple comparisons across time by considering the continuity of adjacent time points. Parameters were set to default for temporal clustering and at each time point, a one-sample t-test was performed against its respective null hypothesis (AUC > 0.5 and ***β*** > 0). Samples exceeding a cluster-forming threshold of p < 0.05 were grouped into temporal clusters based on adjacency. Cluster-level statistics were defined as the sum of t-values within each cluster, and a null distribution was constructed using 5000 permutations with random sign flipping. For reporting purposes, we additionally summarize each significant cluster using the range minimum and the maximum t-value observed within the cluster.

## RESULTS

### Behavioral Shifts from Exploration to Exploitation

Following the introduction of novel stimuli, participants initially favored exploratory over exploitative choices (**Figure 1.B**). We quantified this behavior using POMDP derived estimates of IEV and exploration BONUS. Trials were classified as exploratory when choices were driven by a positive exploration bonus despite relatively low expected value, and as exploitative when choices reflected high expected value in the absence of an exploration bonus. Consistent with an initial bias toward uncertainty-driven exploration, behavior at the onset of the novel phase favored exploratory choices (M = 0.58, SEM = 0.03) over exploitation (M = 0.41, SEM = 0.03). Across subsequent trials, this balance progressively shifted, with exploration decreasing and exploitation increasing as the value of the novel option became established. By the end of the novel phase, choices were predominantly exploitative (M = 0.68, SEM = 0.02), accompanied by a corresponding reduction in exploratory behavior (M = 0.31, SEM = 0.02), (**Figure 1.B**). A binomial general linear model was used to analyze choice dynamics, predicting the likelihood of an exploratory vs. exploitative choice as a function of trials since novel insertion. Consistent with the observed shift from exploration to exploitation, the probability of exploratory choices declined significantly across trials (β = −0.20, SEM = 0.03, t = −7.94, p < .001; **Figure 1.B**), while the probability of exploitative choices increased in parallel (β = 0.19, SEM = 0.03, t = 7.94, p < .001; **Figure 1.B**). Overall, these results indicate that participants initially favored exploration when encountering a novel option, but progressively shifted toward exploiting the most rewarding alternative as experience accumulated.

### Spatiotemporal Dynamics of Evoked MEG Responses

Cue-locked activity in sensor space revealed a dynamic, temporally structured response, with RMS amplitude rising rapidly at ∼100–150ms and peaking around ∼200–300ms (**Figure 2.A,C**), primarily over posterior sensors and localizing to visual cortex (**Figure 2.E**), consistent with early perceptual processing of the choice set. This initial burst was followed by a clear posterior-to-anterior propagation, reflecting a feedforward transition to higher-order associative networks by ∼200–300ms. Within these networks, a hierarchical pattern emerged in which regions such as the intraparietal sulcus and inferior frontal cortex showed early but transient engagement (**Supplementary Figure 1**), while a core rostral prefrontal network, including lateral frontopolar cortex (FPC), ventromedial prefrontal cortex (vmPFC), and orbitofrontal cortex (OFC), became active shortly after the visual peak and remained sustained throughout the ∼300–600ms pre-decision window. Feedback-locked activity engaged a largely overlapping cortical network but with a distinct temporal profile, including an early visual response at ∼150–250ms (**Figure 2.B,D**), alongside sustained activation of higher-order associative regions beyond ∼500ms (**Figure 2F**; **Supplementary Figure 1**). In short, both cue and feedback-locked MEG responses demonstrate a combination of transient visual cortical responses and sustained engagement of parietal, temporal, and frontal association areas. With this neurophysiological baseline of the MEG responses during our task clearly established, we next turn to model-based decoding of the MEG signal during these events to resolve the specific sequence of cortical computations that shape explore-exploit decision policies and their evaluation in real time.

**Figure 2.**
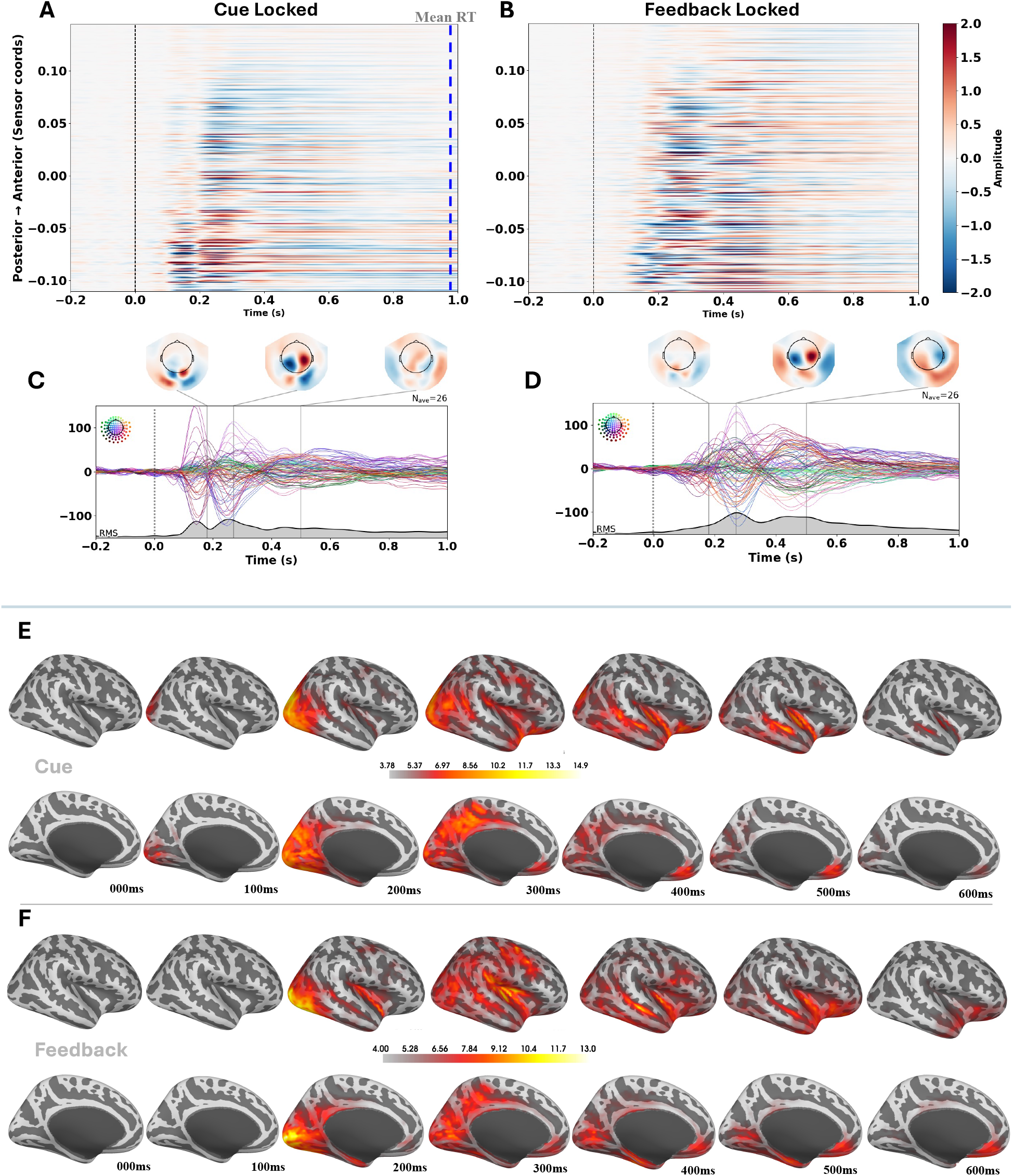
Sensor and Source level spatiotemporal evolution of evoked responses. (A–B) Heatmaps showing normalized sensor (magnetometers and gradiometers) amplitudes for (A) cue-locked and (B) feedback-locked periods. Sensors are ordered along the posterior-to-anterior axis to illustrate the characteristic signal propagation over time. (C) Grand-average waveforms across all sensors (bottom) and representative topographies (top) showing a focal, transient peak in posterior sensors at cue. (D) Grand-average waveforms and topographies following feedback onset. Compared to cue, feedback processing exhibits more sustained activity and a broader spatial distribution extending into anterior sensor groups. (C– D, grey shading) Root mean square (RMS) of the signal across all sensors, highlighting the global field power over time. (E) Posterior-to-anterior propagation during cue processing. Source estimates (0–600 ms) show early occipital engagement (∼100 ms) followed by a sweep of activity toward associative parietal and frontal regions. (F) Feedback-locked activity shows more persistent and spatially broad cortical responses (300–600 ms) compared to the cue-locked cascade.

### Spatiotemporal Decoding of Explore-Exploit Decision and Outcome Variables

To determine what computations drive these spatiotemporal dynamics, we decoded task variables directly from the MEG activity. At cue, explore versus exploit choices were reliably classified relatively early and remained intermittently decodable up until choice execution (∼200ms–1000ms; mean AUC ∼0.55–0.58; sensor t = [1.767, 5.791], p < 0.01; source t = [1.929, 3.431], p < 0.05; **Figure 3.A.i**). Since our binary choice decoder labels inherently confound the distinct computational parameters that drive them (e.g. value versus information seeking), we sought to isolate these signals by regressing trial-wise decoder decision values from test trials onto the orthogonalized latent variables. This revealed a significant, sustained divergence in the encoding of immediate expected value (IEV) and exploration value (BONUS) beginning ∼100–150ms after cue onset and extending throughout the decision window (∼800ms) (**Figure 3.A.ii**). At feedback, classifiers robustly discriminated rewarded versus non-rewarded outcomes (mean AUC ∼0.65–0.70; sensor t = [1.785, 11.288], p < 0.001; source t = [1.752, 13.531], p < 0.001), peaking between ∼300–500ms (**Figure 3.B**). Visualizing the spatial progression of these decoder maps (Haufe-coefficients) revealed a highly dynamic, feedforward flow of task-relevant information. The resulting cortical maps demonstrate a sweeping posterior-to-anterior cascade during both decision formation and feedback evaluation (**Figure 3.C-D**).

**Figure 3.**
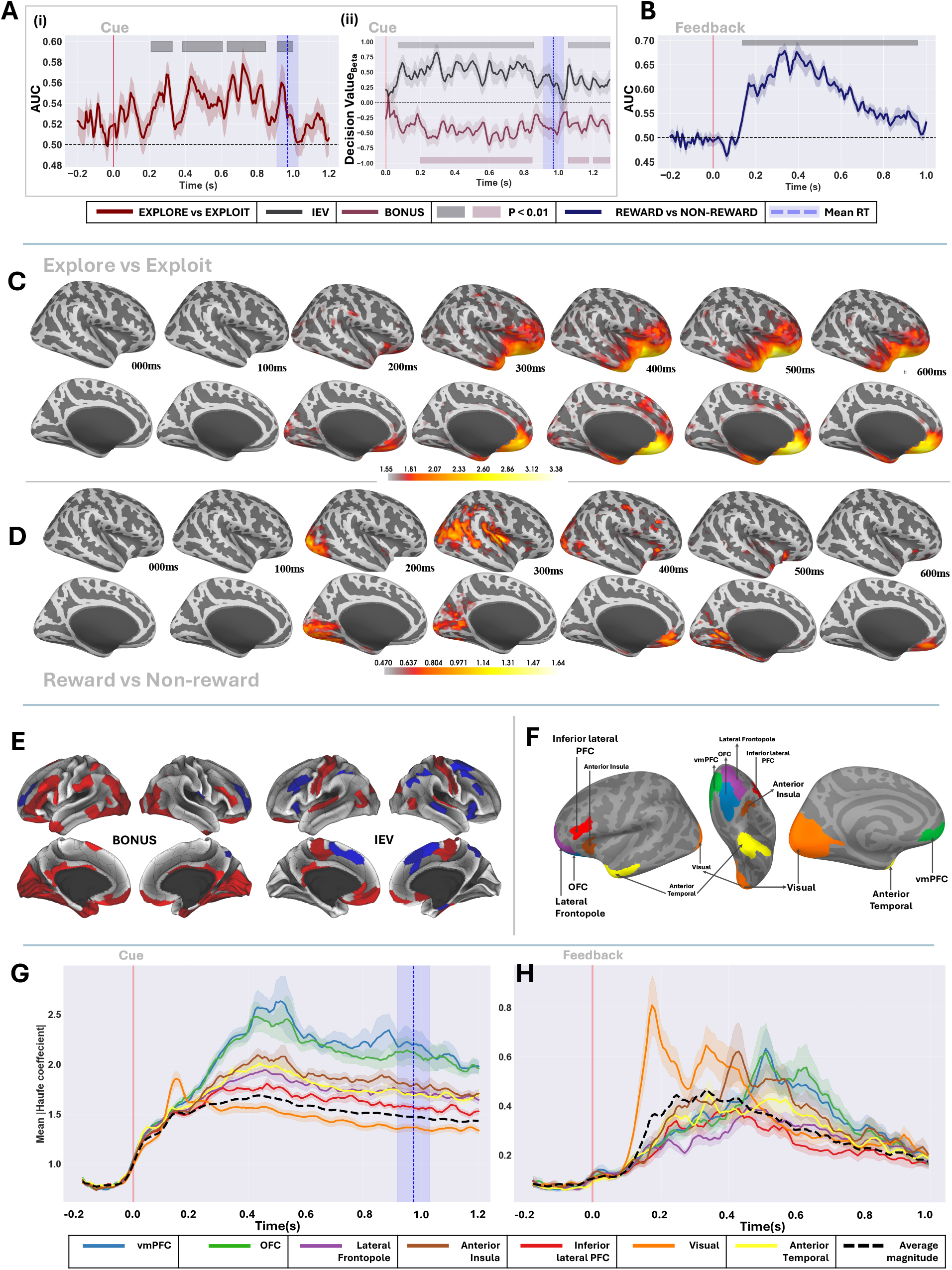
Temporal decoding performance, source-level haufe maps and ROI-level decoding dynamics. (A) (i) Time-resolved decoding performance (AUC) for explore versus exploit classification, time-locked to cue onset. (ii) Decision values from the decoder showing temporal traces of IEV and BONUS. (B) Time-resolved decoding performance (AUC) for reward versus non-reward classification, time-locked to feedback onset. (C) Source-level Haufe activation patterns from explore versus exploit decoding. Maps reflect the magnitude of task-relevant information across cortical sources. (D) Source-level Haufe activation patterns for reward versus non-reward decoding. Maps reflect the magnitude of outcome-related information across cortical sources during outcome evaluation. (E) Cortical regions identified in prior fMRI study as encoding key explore-exploit variables (Hogeveen et al., 2022), used to guide the current ROI definitions. (F) Functional ROIs defined using the Glasser atlas (Glasser et al., 2016; Hogeveen et al., 2022) serving as the primary units for subsequent time-resolved encoding analyses, including prefrontal, parietal, and visual cortices. (G) ROI-averaged Haufe coefficient magnitude for explore versus exploit decoding (cue-locked). Shaded areas represent SEM. Blue dotted line and shaded region represent the mean reaction time +/-95%-CI. (H) ROI-averaged Haufe coefficient magnitude for reward versus non-reward decoding (feedback-locked). Colored lines represent different cortical regions, with the dashed line indicating the average magnitude across ROIs.

To localize the precise cortical drivers of these computational signals at cue and feedback time, we examined Haufe-transformed activation patterns from the source-space decoders within a priori regions of interest (ROIs identified via (Hogeveen et al., 2022); **Figure 3.E-H**). During the cue phase, decoding-related activity was initially driven by a transient visual cortex response (∼150–200ms). Following this early sensory sweep, higher-order regions, specifically the vmPFC, OFC, lateral frontopolar (FPC), and anterior temporal cortex, emerged as the dominant contributors. These regions exhibited sustained engagement from ∼250ms onward, peaking between ∼400–500ms (**Figure 3.G**). A temporally shifted but anatomically analogous cascade occurred during outcome evaluation (**Figure 3.H**). An early visual response (∼150–250ms) was rapidly followed by robust, sustained recruitment of the same prefrontal and anterior temporal network from ∼250ms onward. This overlapping spatiotemporal progression confirms a shared cortical substrate that transitions from encoding pre-choice decision variables to evaluating post-choice outcomes.

#### Sustained Prefrontal Encoding of Empirical Value at Feedback

To determine whether outcome-locked neural representations encode behaviorally relevant value signals, trial-wise decision values (***z***_***o***_) at test trials (**Figure 3.D)**, were extracted from the source-space decoders at feedback. Plotting these decision values separately for reward and non-reward trials (**Figure 4.A**) revealed a clear temporal divergence following feedback onset. These decision values were regressed onto the latent computational variables (IEV and BONUS), revealing a sustained representation of empirical value over an extended temporal window regardless of the specific trial outcome (i.e., Reward or Non-Reward; (**Figure 4.B**)). Notably, significant clusters (*p* < 0.01) indicated a reliable representation of IEV that emerges early (∼150-250ms) and remains intermittently significant over an extended temporal window up to ∼900ms post-feedback. In contrast, BONUS showed no consistent significant encoding across the feedback period. Aligning these encoding profiles with the source-resolved decoding data demonstrates that the vmPFC and OFC maintain a sustained tracking state during this same post-outcome window. Together, this indicates that post-choice prefrontal representations are strictly dedicated to tracking empirical choice value, providing a necessary bridge to update valuations and shape future choice policies.

**Figure 4.**
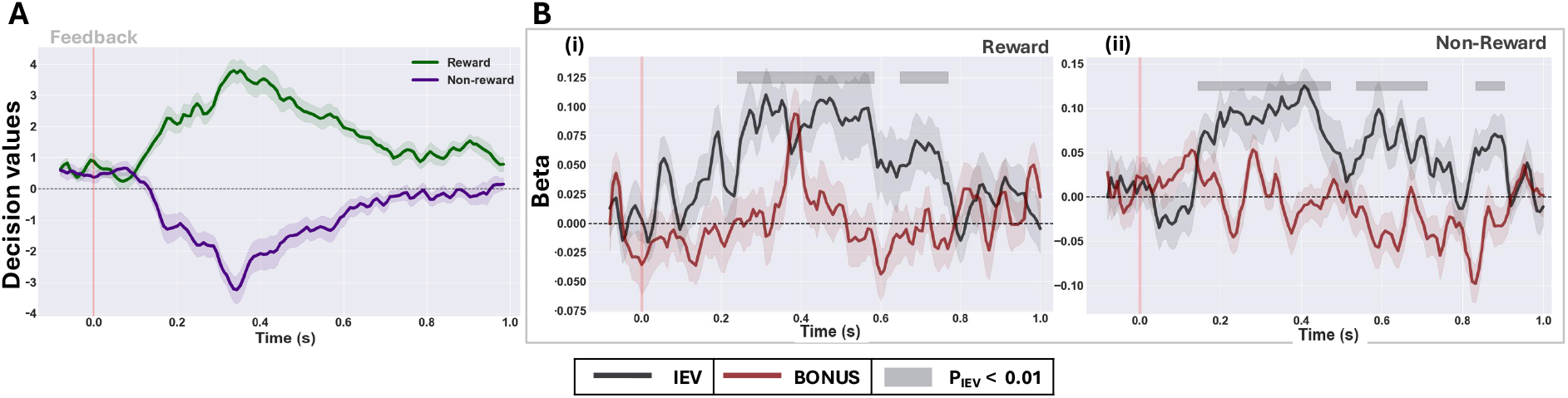
Temporal dynamics and computational encoding of decision values at feedback. (A) Time-resolved decoder decision values plotted separately for reward (green) and non-reward (purple) trials, showing a clear temporal divergence following feedback onset. (B) (i) Reward Trials: Neural encoding of decision values tracks IEV when choice resulted in a reward. (ii) Non-Reward Trials: Similar encoding profile emerges for IEV when choice resulted in no reward.

#### Frontopolar Cortex Initiates Strategic Exploration

While robust neural responses to reward versus non-reward feedback and encoding of empirical value in vmPFC and OFC points to a role for these regions in choice valuation and learning to exploit, this encoding cannot account for the drive to initiate exploit-to-explore shifts during novelty-driven exploration. To isolate the precise temporal onset of strategic deviation from the baseline exploit state across prefrontal circuitry, a posthoc mass univariate regression model was applied to the vmPFC, OFC, and FPC ROIs that emerged in our explore-exploit decoding model at the cue-locked decision window. The activity profiles from this regression model reveal stark explore-exploit functional dissociation across the rostral prefrontal cortex (**Figure 5.A-C**). Within the FPC, exploratory trials significantly deviated from the exploitative baseline early in the decision phase. This divergence emerged as a broad, sustained significant cluster (*p* < 0.01, *t* = [-0.23, 9.96]; **Figure 5.D**) beginning at ∼580ms and remaining continuously significant throughout choice execution, displaying an enhancement in beta magnitude that peaked well in advance of the average reaction time (M = 970ms, 95% CI [910, 1029]).

**Figure 5.**
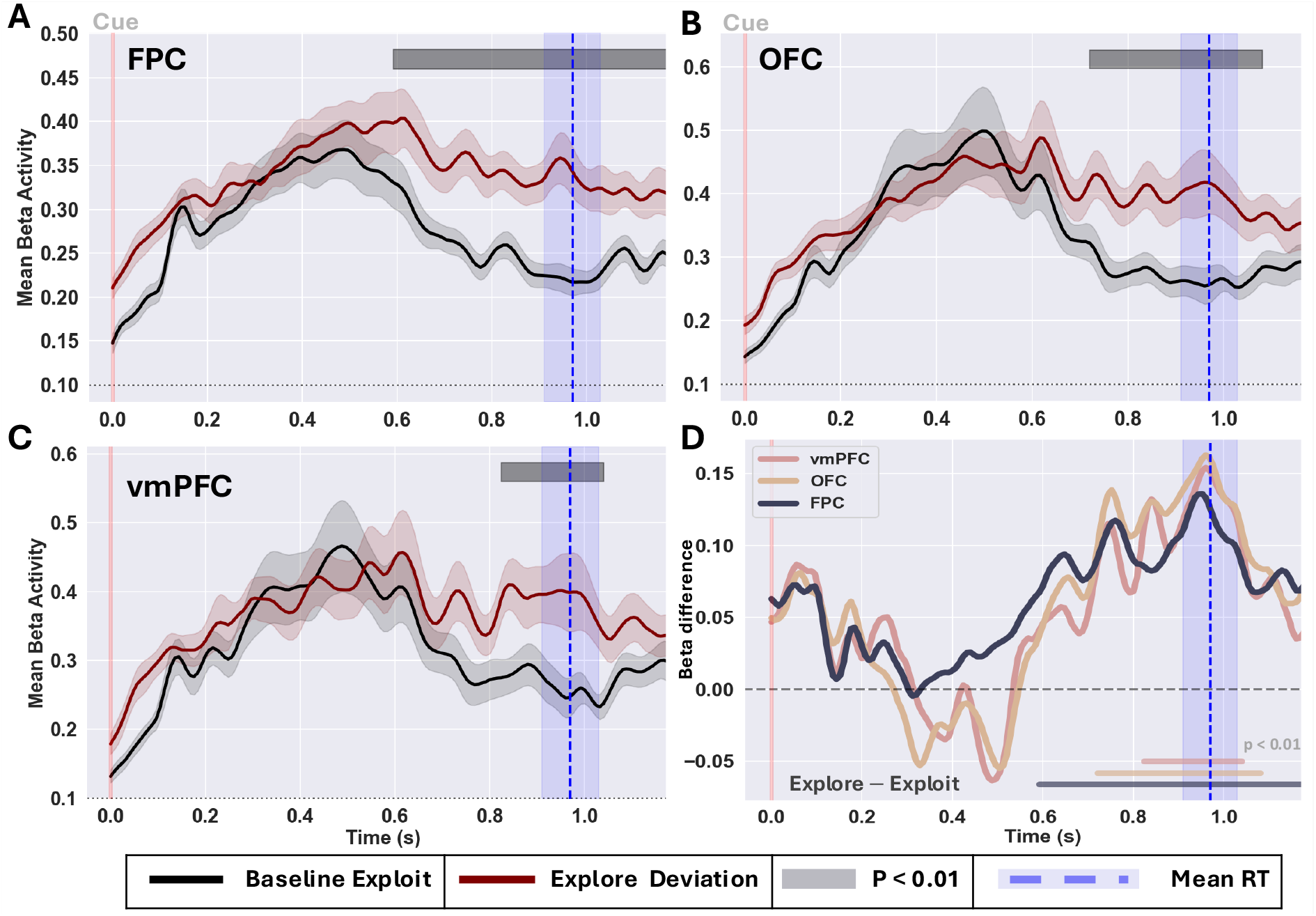
Differential neural recruitment during exploration across prefrontal ROIs from baseline exploitation. (A) **FPC:** Time-resolved beta magnitude demonstrating early, sustained neural divergence from the exploitative baseline during exploration trials. (B) **OFC**: Delayed, transient neural divergence from the baseline exploit trace just prior to choice execution. (C) **vmPFC:** Late-emerging neural divergence restricted to the pre-action window preceding the response. (D) Mean divergence trace highlighting early strategic initiation of exploration in the FPC well ahead of the mean reaction time.

Unlike FPC, neural trajectories within the vmPFC and OFC remained much more tightly coupled to the baseline exploit trace. Rather than a sustained functional shift, both the vmPFC (p < 0.01, t = [-0.85, 6.98]) and the OFC (p < 0.01, t = [-1.03, 7.69]) exhibited brief periods of significant divergence (**Figure 5.D**). These brief shifts occurred preceding and overlapping with mean reaction time (∼700–1000 ms). This temporal dissociation confirms that the FPC initiates and maintains the primary strategic shift away from exploitation, while the vmPFC and OFC only transiently modulate their baseline tracking prior to action execution.

## DISCUSSION

Adaptive decision-making requires organisms to flexibly decide when to exploit familiar rewards, and when to explore novel opportunities that may be more beneficial in the long term (Kaelbling et al., 1996; Sutton & Barto, 2018). A large body of fMRI research has mapped the spatial architecture of computations underlying explore-exploit decision-making (Kobayashi & Kable, 2024; Sazhin et al., 2025; Wyatt et al., 2024). This work has found that frontopolar, ventromedial prefrontal, parietal, and cingulo-opercular regions are involved in, for example, tracking chosen and unchosen action values, and estimating the degree of uncertainty in the choice environment to decide when exploration is advantageous (Badre et al., 2012; Blanchard & Gershman, 2018; Boorman et al., 2009; Cockburn et al., 2021; Daw et al., 2006; Hogeveen et al., 2022; Tomov et al., 2020). However, while the BOLD signal excels at localizing these valuation and control hubs, its temporal constraints have obscured the real-time, algorithmic implementation of explore-exploit decision policies. Magnetoencephalography (MEG) offers an ideal methodological bridge to resolve this limitation. MEG merges the spatial precision necessary to map distributed cortical networks with the millisecond sensitivity required to capture discrete neural computations that are traditionally isolated via EEG (Baillet, 2017; Cavanagh, 2019). In the present study, we leveraged MEG to decode distinct decision states, successfully classifying decision events driven by high latent exploration value (BONUS) from those driven by high exploitation value (IEV). The current findings move beyond static mapping of prefrontal decision-making hubs, demonstrating instead that explore-exploit decision policies emerge from a dynamic, hierarchical cascade that functionally dissociates pre-decision goal formulation from post-decision credit assignment and value updating.

Our observation that the choice to explore or exploit is decodable from visual regions very soon after a choice set is presented suggests there is a feed-forward, visual novelty-based processing that initiates the decision to explore rather than an isolated, prefrontal cortex value comparison mechanism (cf., (Jaegle et al., 2019); **Figure 3.C,G**). This is in line with our recent finding that a novelty-sensitive event-related potential (P3a) is associated with the subsequent drive to explore visually-salient cues (Campbell et al., 2023). Notably, this did not imply that prefrontal decision-making hubs were uninvolved in the formulation of an exploratory versus exploitative choice policy. Instead, there was a clear signal propagation to anterior cortical regions well in advance of choice execution, which revealed a functional dissociation in rostral prefrontal regions. FPC actively initiated the strategic shift toward exploration, exhibiting a sustained divergence from the baseline exploitative state well in advance of choice execution (**Figure 5.A**). In contrast to the FPC, early neural responses within vmPFC and OFC remained statistically matched to the exploitative condition early in the cue epoch, instead demonstrating a delayed, transient exploratory trial modulation just prior to choice execution (**Figure 5.B-C**). This temporal dissociation aligns with the theoretical necessity for a distributed brain network to integrate relative and total uncertainty estimates (Tomov et al., 2020), before goal-directed resolution can be achieved via specialized prefrontal hubs necessary for adaptive decision-making (Zajkowski et al., 2017; Mansouri et al., 2017). Rather than a monolithic prefrontal computation, the human cortex navigates the explore-exploit dilemma through an early, sustained frontopolar initiation of an exploratory choice policy, that may drive a later immediate → future expected value “override” within vmPFC and OFC (**Figure 5.D**).

Following choice execution, the prefrontal network functionally reconfigured. Rather than continuing to track the latent exploration value of choice policies (BONUS), neural decoding shifted exclusively to encoding empirical value (IEV) at feedback time. Reward versus nonreward was decodable ∼100ms after outcome onset; the classifier output displayed a longer, more sustained positive value for reward events and a briefer, less marked deflection for nonreward events (**Figure 4.A**). Time-resolved regression of this feedback-locked neural activity revealed that the vmPFC and OFC transitioned into a stable, sustained representational state dedicated entirely to the immediate expected value (IEV; **Figure 4.B**) of the chosen option. Notably, the spatiotemporal profile of this post-decision feedback decoding showed marked, prolonged activity (e.g., ≥400ms) in the vmPFC (**Figure 3.D,H**). This directly aligns with recent findings on the canonical reward positivity (RewP), an electrophysiological signature of reward processing that has a robust source generator in the vmPFC and is blunted in patients with clinical anhedonia and reduced behavioral reward sensitivity (Pirrung et al., 2025). These findings are also in line with the purported role of vmPFC in credit assignment, selectively tracking IEV to update the current goal state and calibrate reward expectations to shape future decisions (Akaishi et al., 2016; Noonan et al., 2017). Collectively, these findings suggest the neural response to reward in vmPFC tracks and updates expected value, playing a critical role in shaping motivated behavior and subjective affect.

We observed a dissociation whereby feedback-locked neural decoding was aligned with IEV but not the value of exploration (BONUS). The absence of cortical BONUS encoding at outcome evaluation must be contextualized alongside the physiological limitations of the MEG signal. MEG is largely blind to deep subcortical sources. Nonhuman primate electrophysiological studies have clearly demonstrated that BONUS encoding during feedback monitoring can be observed in the amygdala and striatum (Costa et al., 2019), core ventral subcortical motivational brain regions involved in encoding goal or state values to drive reinforcement learning (Averbeck & Murray, 2020). Conversely, cortical representations of the latent value of exploration ramp up early and are strongest during the pre-decision portion of the Bandit task (Costa & Averbeck, 2020; Tang et al., 2022). The absence of feedback-locked MEG decodability of trial-by-trial BONUS values may, therefore, be unsurprising in light of the subcortical imaging limitations of this modality.

Ultimately, operationalizing explore-exploit behavior through a formal POMDP model confirms that human decision-makers positively weigh both immediate empirical value and future information gain. The insertion of novel options reliably triggered uncertainty-driven exploration, which systematically transitioned to exploitation as empirical value was established. By classifying these discrete computational states within millisecond-resolved MEG data, we provide direct evidence that navigating the tradeoff between the known and the unknown relies on an intricately timed, hierarchically organized cortical sequence. This sequence comprises an early visual novelty signal that cues the frontopolar initiation of exploratory policies, a delayed ventromedial and orbitofrontal valuation override prior to exploratory action, and a stable post-decision prefrontal encoding of empirical value to optimize future decision-making.

## Supporting information

Supplemental ROI Evoked activity

Supplemental Regression model

## ACKNOWLEDGMENTS

This work was supported by the National Science Foundation award number #2237795, as well as the National Institutes of Health (via the National Institute of General Medical Sciences (P30GM122734) and the National Institute on Alcohol Abuse and Alcoholism award number (R01AA030283)). We greatly appreciate the help of Drs. Teagan Mullins and Kimberly Kundert-Obando, and Ms. Elizabeth Eversole for their help with data collection for this study.

## Footnotes

1 https://www.mrn.org/collaborate/mind-input-device

