## Supplemental ROI Evoked activity for "Spatiotemporal Decoding of Explore-Exploit Decisions in the Human Brain"

### Supplementary

#### *ROI-level evoked activity*

To further examine region-level source activity, cortical areas were grouped into anatomically and functionally defined regions of interest (ROIs; **Figure 1 Supplementary**) based on the Glasser atlas. The ROIs included early visual cortex, parietal regions, anterior temporal areas, and prefrontal subdivisions such as orbitofrontal cortex (OFC), ventromedial prefrontal cortex (vmPFC), and lateral frontopolar cortex (FPC). This organization was intended to capture the cortical processing hierarchy, spanning early sensory encoding, associative processing, and higher-order evaluative and decision-related functions. Cue-locked responses (**Figure 1 Supplementary. A, C**) were initially dominated by a sharp increase in visual cortex activity peaking around 150–200ms. After excluding visual cortex, a clearer progression emerged: parietal regions, particularly the intraparietal sulcus, and anterior temporal cortex showed early increases (~150–250ms), followed by sustained activation in prefrontal regions, including OFC and vmPFC, peaking around 300–500ms. A similar pattern was observed for feedback-locked responses. Including the visual cortex revealed a prominent early visual peak around 200 ms (**Figure 1 Supplementary. B, D**), whereas excluding it highlighted sustained engagement of higher-order regions. In particular, vmPFC and OFC showed robust and prolonged activation (~300–700 ms), accompanied by contributions from parietal and anterior temporal cortices. Compared with cue processing, feedback processing elicited stronger and more sustained activation across associative and prefrontal regions.

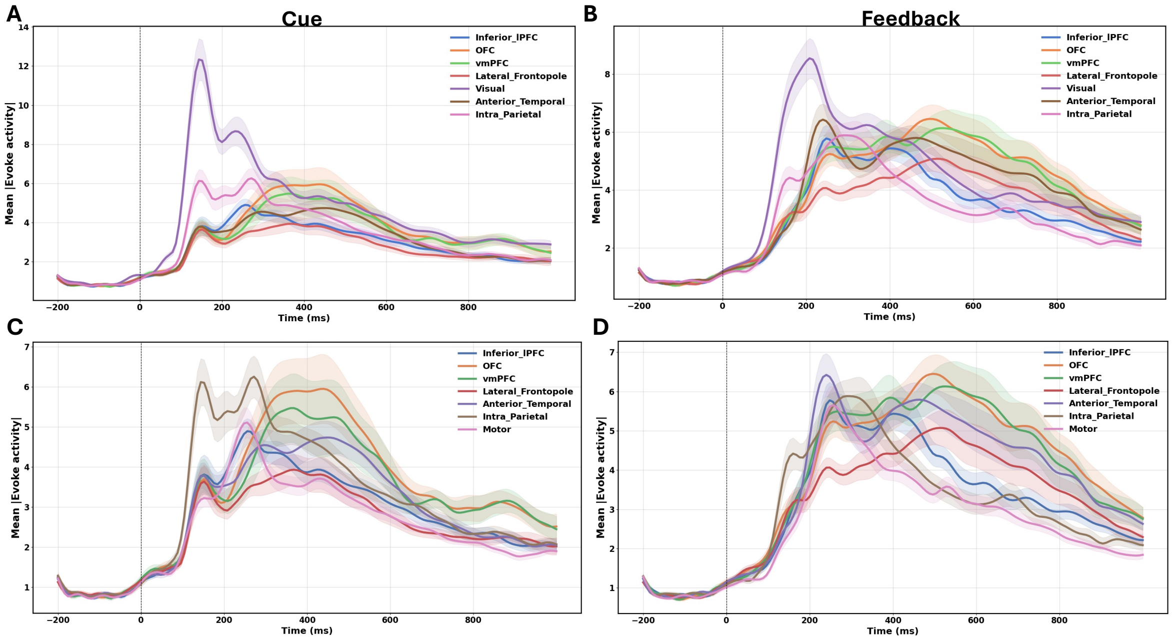


**Figure 1 Supplementary.** ROI-level evoked activity for cue and feedback-locked responses. Mean evoked activity (± SEM) extracted from functionally defined cortical ROIs based on the Glasser atlas (Glasser et al., 2016), shown for cue-locked (A, C) and feedback-locked (B, D) epochs. Panels A and B include all ROIs, with the visual cortex displaying the dominant early response in both conditions. Panels C and D replot the same data with the visual ROI excluded, revealing the relative temporal dynamics of prefrontal (OFC, vmPFC, lateral frontopolar), parietal (intra-parietal), anterior temporal, and motor regions. Shaded regions represent ± SEM across participants. Dashed vertical lines indicate stimulus onset (t = 0 ms).

Taken together, both cue and feedback processing involve coordinated activity across a distributed network, but with distinct temporal profiles. Cue-related activity transitions relatively quickly from posterior to anterior regions, whereas feedback-related processing is characterized by prolonged engagement of prefrontal and associative cortices. These ROI-level findings provide a clearer view of large-scale cortical dynamics and further support the interpretation derived from sensor- and source-level analyses.
