## Supplemental Regression model for "Spatiotemporal Decoding of Explore-Exploit Decisions in the Human Brain"

### Supplementary

#### 1 Mass-Univariate Linear Regression of Outcome-Locked Value Representations

To determine whether outcome-locked neural representations dynamically encode behaviorally relevant value signals, we evaluated the post-feedback timecourse of trial-wise decision values using a comprehensive mass-univariate linear regression model. Rather than fitting separate, stratified models for reward and non-reward trials—which can slice the data arbitrarily, introduce selection bias, and distort parameter estimates—we collapsed all trial types into a single unified estimation framework.

Specifically, the time-resolved decoder decision values $z_o(t)$ at each time point $t$ following outcome onset were regressed onto orthogonalized latent computational variables representing the Expected Value of the chosen option ($IEV_{\text{orth}}$), Reward Prediction Error ($RPE_{\text{orth}}$), and exploration bonus ($Bonus_{\text{orth}}$). To independently map how these latent variables are modulated by the trial's actual feedback while preserving statistical power and avoiding estimation bias, we interacted each computational regressor with condition-specific identity (indicator) functions, $\mathbb{I}(\text{Rew})$ and $\mathbb{I}(\text{NoRew})$. These indicator functions act as binary switches—evaluating to 1 for trials matching the specific outcome condition and 0 otherwise—thereby partitioning the variance cleanly between reward and non-reward trials within the same regression matrix. The complete model is specified as follows:

$$\begin{aligned} z_o(t) = \beta_0(t) &+ \beta_1(t) \cdot IEV_{\text{orth}} \cdot \mathbb{I}(\text{Rew}) + \beta_2(t) \cdot IEV_{\text{orth}} \cdot \mathbb{I}(\text{NoRew}) \\ &+ \beta_3(t) \cdot RPE_{\text{orth}} \cdot \mathbb{I}(\text{Rew}) + \beta_4(t) \cdot RPE_{\text{orth}} \cdot \mathbb{I}(\text{NoRew}) \\ &+ \beta_5(t) \cdot Bonus_{\text{orth}} \cdot \mathbb{I}(\text{Rew}) + \beta_6(t) \cdot Bonus_{\text{orth}} \cdot \mathbb{I}(\text{NoRew}) + \varepsilon_i(t) \end{aligned}$$
